# Nutrient dominance governs the assembly of microbial communities in mixed nutrient environments

**DOI:** 10.1101/2020.08.06.239897

**Authors:** Sylvie Estrela, Alicia Sanchez-Gorostiaga, Jean C.C. Vila, Alvaro Sanchez

## Abstract

A major open question in microbial community ecology is whether we can predict how the components of a diet collectively determine the taxonomic composition of microbial communities. Motivated by this challenge, we investigate whether communities assembled in pairs of nutrients can be predicted from those assembled in every single nutrient alone. We first find that although the null, naturally additive model generally predicts well the family-level community composition, there exist systematic deviations from the additive predictions that reflect generic patterns of nutrient dominance at the family-level. Pairs of more similar nutrients (e.g. two sugars) are on average more additive than pairs of more dissimilar nutrients (one sugar-one organic acid). Second, a simple dominance rule emerges: sugars generally dominate organic acids. These findings may be explained by family-level asymmetries in nutrient benefits. Overall, our results suggest that regularities in how nutrients interact may help predict communities responses to dietary changes.

## Introduction

Understanding how the components of a complex biological system combine to produce the system’s properties and functions is a fundamental question in biology. Answering this question is central to solving many fundamental and applied problems, such as how multiple genes combine to give rise to complex traits [1,2], how multiple drugs affect the evolution of resistance in bacteria and cancer cells [3,4], how multiple environmental stressors affect bacterial physiology [5], or how multiple species affect the function of a microbial consortium [6–8].

In microbial population biology, a major related open question is whether we can predict how the components of a diet collectively determine the taxonomic and functional composition of microbial communities. Faith and co-workers tackled this question using a defined gut microbial community and a host diet with varying combinations of four macronutrients [9]. This study found that community composition in combinatorial diets could be predicted from communities assembled in separate nutrients using an additive linear model. Given the presence of a host and its own possible interactions with the nutrients and resident species, it is not immediately clear whether such additivity is directly mediated by interactions between the community members and the supplied nutrients, or whether it is mediated by the host, for instance by producing additional nutrients, or through potential interactions between its immune system and the community members. More recently, Enke et al found evidence that marine enrichment communities assembled in mixes of two different polysaccharides could be explained as a linear combination of the communities assembled in each polysaccharide in isolation [10].

Despite the important insights provided by both of these studies, we do not yet have a general quantitative understanding of how specific nutrients combine together to shape the composition of self-assembled communities [11]. Motivated by this challenge, here we set out to systematically investigate whether the assembly of enrichment microbial communities in a collection of defined nutrient mixes could be predicted from the communities that assembled in each of the single nutrients in isolation.

## Results

### A null expectation for community assembly in mixed nutrient environments

To investigate whether communities assembled in pairs of nutrients can be predicted from those assembled in every single nutrient alone, we must first develop a quantitative null model that predicts community composition in a mixed nutrient environment in the case where each nutrient recruits species independently. Any deviation between the null model prediction and the observed (measured) composition reveals that nutrients are not acting independently, but rather “interact” to shape community composition. This definition of an interaction as a deviation from a null model that assumes independent effects is commonplace in systems-level biology [12,13].

In order to formulate the null expectation for independently acting nutrients, let us consider a simple environment consisting of two unconnected demes where two bacterial species, A and B, can grow together. The first deme contains a single growth limiting nutrient (nutrient 1), while the second deme contains a different single limiting nutrient (nutrient 2) (**Fig. 1A**). In this scenario, each nutrient influences the abundance of species A and B independently: the microbes growing on nutrient 1 do not have access to nutrient 2 and vice versa. Let’s denote the abundance of species A in demes 1 and 2 by *n*_*A,1*_ and *n*_*A,2*_, and the abundance of species B as *n*_*B,1*_ and *n*_*B,2*_, respectively. If we now consider the two-deme environment as a whole, the abundance of species A is the sum of its abundance in each deme *n*_*A,12*_ = *n*_*A,1*_ + *n*_*A,2*_ (likewise, for species B *n*_*B,12*_ = *n*_*B,1*_ + *n*_*B,2*_). This example illustrates that in the scenario when two limiting nutrients act independently, each of them recruits species just as if the other nutrient were not there. In such case, the abundance of each species in a nutrient mix is the sum of what we would find in the single nutrient habitats. Note that the lack of nutrient interactions does not mean that species do not interact with each other, but rather that whatever ecological or metabolic interaction they may have (e.g., competition for nutrients, cross-feeding, growth inhibition by toxins), such interaction is not affected by mixing nutrients.

**Fig. 1.**
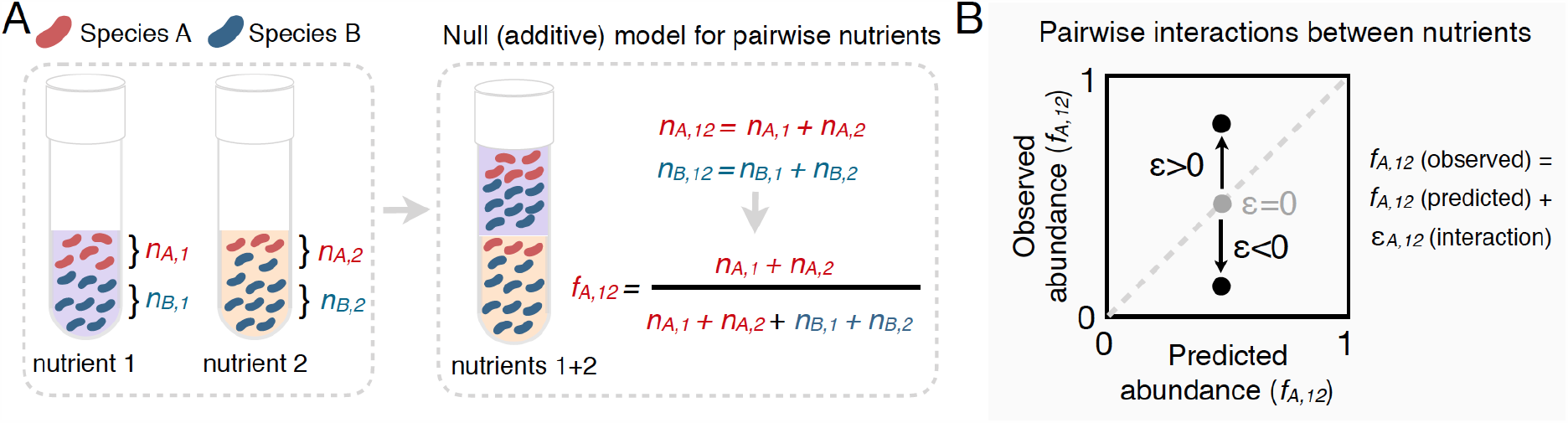
Predicting community composition in mixed nutrient environments. **(A)** Community composition in a single nutrient (nutrient 1 or nutrient 2) *vs* a mixture of nutrients (nutrient 1 + nutrient 2). Assuming that nutrients act independently, the null model predicts that the abundance of each species in the mixture is the sum of its abundance in the single nutrients (i.e. additive). **(B)** Plotting the experimentally measured (observed) relative abundance in the mixed carbon sources against its predicted (from null model) relative abundance reveals the presence or absence of interactions. Any deviation from the identity line (predicted=observed) is the interaction effect (*ε*). When *ε* = 0, there is no interaction between nutrients. When *ε* is non-zero, community composition is affected by nutrient interactions. If *ε*>0, the null model underestimates the abundance. If *ε*<0, the model overestimates the abundance.

Under the null model, the relative abundance of species *i* in a mix of nutrients 1 and 2 can be written as *f*_*i,12*_(null) = *w*_*1*_ *f*_*i,1*_ + *w*_*2*_ *f*_*i,2*_ where *f*_*i,1*_ and *f*_*i,2*_ are the relative abundances of *i* in nutrient 1 and 2, respectively, and *w*_*1*_ and *w*_*2*_ are the relative number of cells in nutrients 1 and 2 (Methods). Any quantitative difference between the null model prediction and the observed composition quantifies an “interaction” between nutrients. Accounting for the presence of such interactions, the model can be re-written as *f*_*i,12*_ = *f*_*i,12*_(null) + *ε*_*i,12*_ where *ε*_*i,12*_ represents the interaction between nutrients 1 and 2 (**Fig. 1B**).

### Experimental system

Equipped with this null model, we can now ask to what extent the nutrients recruit species independently in mixed environments. To address this question, we followed a similar enrichment community approach to the one we have used in previous work for studying the self-assembly of replicate microbial communities in a single carbon source [14,15] (Methods, **Fig. 2A**). Briefly, habitats were initially inoculated from two different soil inocula. Communities were then grown in synthetic (M9) minimal media supplemented with either a single carbon source or a mixture of two carbon sources, and serially passaged to fresh medium every 48h for a total of 10 transfers (dilution factor=125×) (**Fig. 2A**). The carbon source pairs consisted of a focal carbon source mixed 1:1 with one of eight additional carbon sources. We previously found that stable multi-species communities routinely assemble in a single carbon source (which is limiting under our conditions), and they converge at the family level in a manner that is largely governed by the carbon source supplied, while the genus or lower level composition is highly variable [14]. We chose glucose as the focal carbon source because we have previously carried out multiple assembly experiments in this nutrient [14,15]. As the additional carbon sources, we chose cellobiose, fructose, ribose, and glycerol (i.e. a pentose, a hexose, a disaccharide and a sugar alcohol) and fumarate, benzoate, glutamine and glycine (two aminoacids and two organic acids). All carbon sources were also used in single carbon source cultures.

**Fig. 2.**
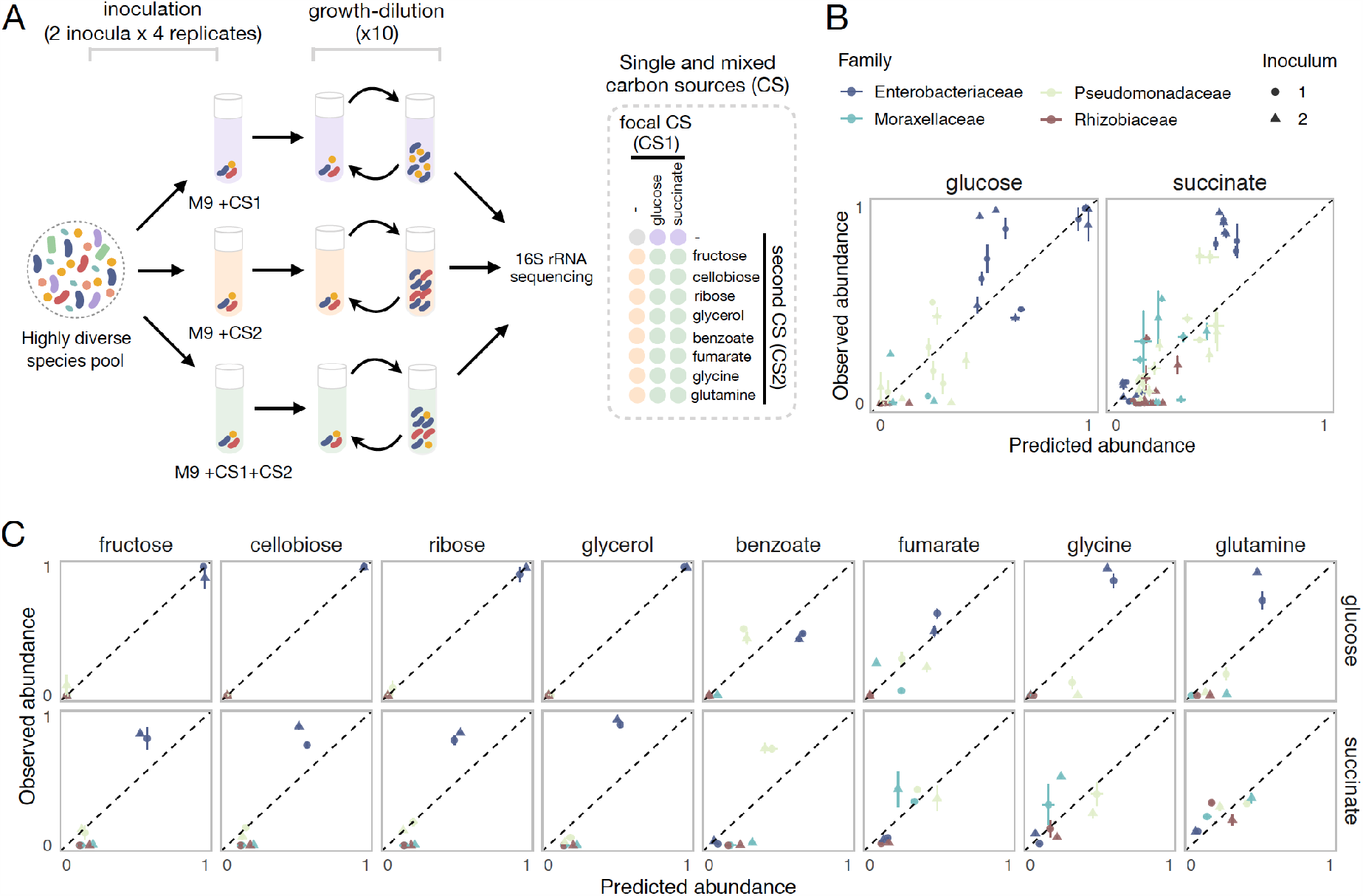
Systematic deviations from the null prediction reveals that some nutrients interact to shape community assembly. **(A)**. Schematic of experimental design. Two different soil samples were inoculated in minimal M9 medium supplemented with either a single carbon source (CS1 or CS2) or a mixture of two carbon sources (CS1 + CS2) (3-4 replicates each). Communities were propagated into fresh media every 48h for 10 transfers, and then sequenced to assess community composition. Carbon source mixtures consisted of a focal carbon source (CS1; glucose or succinate) mixed with a second carbon source (CS2). **(B, C)**. For each pair of carbon sources, we show the experimentally observed and predicted (by the null additive model) relative abundance of each family in the mixture. Any deviation from the identity line (predicted=observed) reveals an interaction effect. Only the four most abundant families are shown. Error bars represent mean ± SE.

Communities assembled in single sugars contained 5 to 24 ESVs, mainly belonging to the Enterobacteriaceae family (∼0.98±0.03), a sugar specialist (**Fig. S1**). In contrast, communities assembled in organic acids exhibited a higher richness (12-36 ESVs), and unlike in sugars, Enterobacteriaceae were generally rare (∼0.06±0.06). Instead, communities were dominated by respirative bacteria mainly belonging to the Pseudomonadaceae (∼0.51±0.25), Moraxellaceae (∼0.18±0.21), and Rhizobiaceae (0.11±0.13) families (**Fig. S1**). Because of the observed family-level convergence across carbon sources, which is consistent with previous studies [14–16], we focus our analysis below on family-level abundance.

### The null model of independently acting nutrients explains a high fraction of the variation observed

To investigate the predictive power of the null (additive) model, we compare the predicted and observed relative abundances of each family for each carbon source pair across all experiments. Our results show that the null model predicts reasonably well the family-level abundances on average (Pearson’s R=0.95 and p<0.001; RMSE=0.073, N=223) (**Fig. 2B, Fig. S2, S3**). To confirm that the strong predictive power of the null model is not an idiosyncrasy of using glucose as the focal carbon source in the pairs, we repeated the same experiment with succinate (an organic acid) as the focal carbon source. Although the correlation between observed and predicted abundance is lower than for glucose, the null additive model is still predictive (Pearson’s R=0.87 and p<0.001; RMSE=0.094; N=257) (**Fig. 2B**).

This result seems to indicate that, at the family level, a simple model that assumes that nutrients act independently can predict community composition in a pair of nutrients (for an analysis of this point at the genus and ESV level, see **Fig. S4**). However, when we looked at this more closely and broke down our results by carbon source and family, we found consistent and systematic deviations from the null model (**Fig. 2C**). For example, across all succinate-sugar pairs, Enterobacteriaceae are significantly more abundant than predicted by the null model (*ε =* 0.347 ± 0.107, Mean±SD; p<0.001, one-sample Student’s t-test, N=32) while both Rhizobiaceae and Moraxellaceae are less abundant than predicted (*ε =* -0.136 ± 0.0339 and *ε =* -0.152 ± 0.0415; p<0.001, one-sample Student’s t-test, N=32) (**Fig. 2C**). The null ‘interaction-free’ model also predicts species abundance better in certain carbon source combinations (e.g. glucose + ribose) than in others (e.g. glucose + glutamine) (**Fig. 2C**). The existence of systematic deviations from the null prediction reveals that some nutrient pairs do not recruit families independently, but instead “interact” with each other to affect the abundance of specific families.

### A simple dominance rule in mixed nutrient environments: sugars generally dominate organic acids

To map the regularities in nutrient interactions observed, we next sought to characterize the nature of these interactions for each carbon source pair and every family. One helpful way of visualizing nutrient interactions is to draw the pairwise abundance landscape for each species and carbon source pair (**Fig. 3A**). For instance, a species could be either more abundant in a pair of nutrients than it is in any of them independently (synergy). Or it could be less abundant than it is in any of the two (antagonism). Dominance is a less extreme interaction which can be visualized by the pushing of a species abundance towards the value observed in one of the two nutrients and away from the average, thus overriding the effect of the second available nutrient (**Fig. 3A**).

**Fig. 3.**
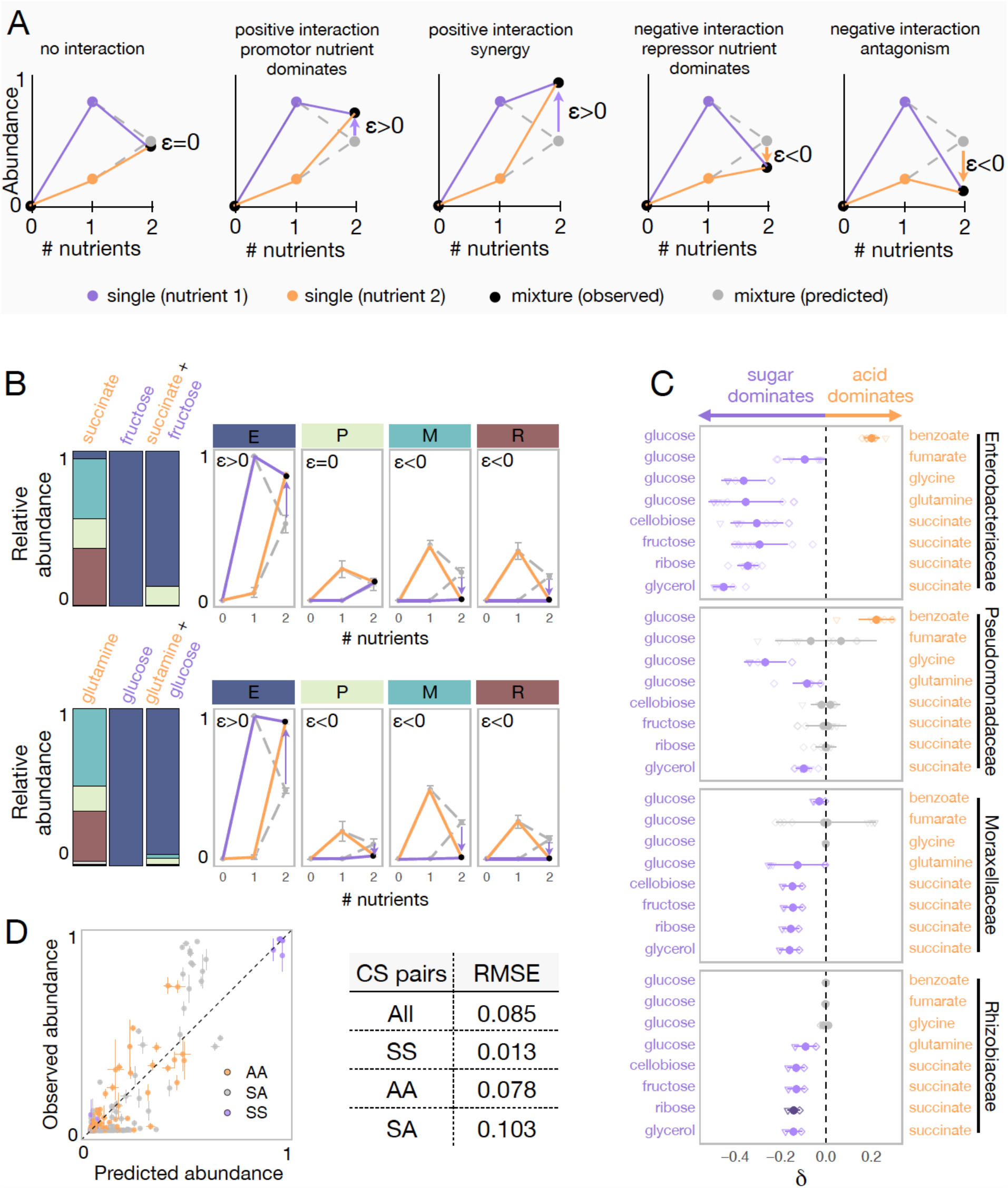
Sugars generally dominate over organic acids. **(A)** Detecting interactions and hierarchies of dominance between nutrients on microbial community composition. Drawing the single and pairwise abundance landscapes for each species allows us to visualise interactions between nutrients. Multiple types of interactions are possible, including dominance, synergy and antagonism. Interactions occur when *ε* is significantly different from 0 (one-sample Student’s t-test, p < 0.05). Synergy (antagonism) occurs when the abundance in the mixture is greater (lower) than the abundances in any of the single nutrients independently (Methods). Dominance occurs when the abundance in the mixture is closer or similar to the abundance in one of the singles. The landscape also allows us to identify which carbon source has a dominating effect within the pair. When ε>0, the growth-promoting nutrient dominates and has an overriding effect in the community composition. In contrast, when ε<0, the growth-repressing nutrient dominates. **(B)** Two examples of nutrient interactions (succinate + fructose and glucose + glutamine) exhibiting sugar dominance. Barplots show a representative replicate from one of the inocula (**Fig. S1, S2**). For instance, the landscape for succinate-fructose shows that fructose overrides the effect of succinate by promoting Enterobacteriaceae (E), and repressing Moraxellaceae (M) and Rhizobiaceae (R) (purple arrows), whereas no interaction effect is observed for Pseudomonadaceae (P). Error bars represent mean ± SD of the four replicates. **(C)** Panel shows dominance index for the eight sugar-acid pairs and the four dominant families. Filled circles show the mean±SD of the 2 inocula x 4 replicates for each pair of nutrients, and open symbols show all 8 independent replicates (different shapes for different inocula). Purple indicates that the sugar dominates while orange indicates that the acid dominates. Lighter purple and orange indicate dominance while darker purple and orange indicate super-dominance (synergy or antagonism). No interaction is shown in gray, and occurs when *ε=*0 (one-sample Student’s t-test, p < 0.05, N=8) or when dominance is undefined because the two inocula exhibit opposite dominant nutrient (in which case *δ* is shown as both -*δ and +δ*). **(D)** Predicted vs observed family-level abundance. For each pair of carbon sources (CS), shown is the experimentally observed and predicted (by the null model) relative abundance of each family in the mixed carbon sources. Any deviation from the identity line (predicted=observed) is the interaction effect. The colours show whether the carbon source pairs are sugar-sugar (SS), acid-acid (AA), or sugar-acid (SA). Error bars represent mean ± SE. Table shows RMSE for each carbon source pair type.

When the interaction is positive (*ε*>0), the dominant nutrient is the one where the family grew to a higher abundance. When the interaction is negative (*ε*<0), the dominant nutrient is the one where the species grew less well. Mathematically, dominance occurs when |*ε*|>0 and *min*(*f*_*i,1*_, *f*_*i,2*_) ≤ *f*_*i,12*_ ≤ *max*(*f*_*i,1*_, *f*_*i,2*_), while synergy and antagonism (forms of super-dominance) occur when |*ε*|>0 and *f*_*i,12*_ > *max*(*f*_*i,1*_, *f*_*i,2*_) and *f*_*i,12*_ < *min*(*f*_*i,1*_, *f*_*i,2*_) respectively (Methods). **Fig. 3B** shows representative examples of dominant carbon source interactions. For instance, Moraxellaceae and Rhizobiaceae grow strongly on succinate, but they are not found in fructose. When fructose is mixed with succinate, both families drop dramatically in abundance, despite their high fitness in succinate alone. Interestingly, however, the dominance of fructose over succinate is not observed for all families: those two nutrients do not interact on Pseudomonadaceae, whose abundance is well predicted by the null model. Using this framework, we then systematically quantified the prevalence of dominance, antagonism and synergy between nutrients for each family **(Fig. S5A)**. While 59% of the nutrient pair combinations exhibited no significant interaction, dominance was by far the most common interaction amongst those that interacted (73%, **Fig. S5A**). It occurred predominantly in the sugar-acid pairs, and to a lesser extent in the acid-acid pairs, and only rarely in the sugar-sugar pairs (**Fig. S5B**). This result strongly suggests that nutrient interactions are not random but do have a specific structure that is conserved at the family-level (**Fig. S5C**).

To systematically characterize and quantify nutrient dominance, we developed a dominance index (*δ*) (Methods). For visualization purposes, the dominance index for the sugar-acid pairs (we will discuss the acid-acid pairs later) is written as *δ*_*i*_ = -|*ε*_*12*_| when the sugar dominates and as *δ*_*i*_ = |*ε*_*12*_| when the acid dominates. If *ε*_*12*_ *=* 0, then *δ*_*i*_ = 0. That is, in the absence of interaction between nutrients, there is no dominance. By plotting the dominance index for each pair of nutrients and each family, we observe a generic pattern of dominance of sugars over acids (**Fig. 3C**). The families Moraxellaceae or Rhizobiaceae are recruited to the community by most organic acids in isolation, but they are not found in most sugar communities. When sugars and organic acids are mixed together, the sugar dominates and both families are at much lower abundances (by ∼6-fold in the case of Moraxellaceae and ∼114-fold in Rhizobiaceae) than expected by the null model, even though the organic acid where they thrived is present in the environment. Consistent with this pattern, we found that pairs of more similar nutrients (a pair of sugars or a pair of organic acids) were significantly better predicted by the null model than mixed organic acid-sugar pairs (**Fig. 3D**). No generic pattern of dominance was observed in the acid-acid mixtures (**Fig. S6**). When we examine interactions and dominance at the genus-level, we find that sugars do not exhibit the same dominance for all genera within the same family (**Fig. S7 and S8**). This result is consistent with the convergence of community structure at the family level (despite substantial variation at lower levels of taxonomy) which we have reported for communities assembled in a single nutrient [14,15]. Together, these results indicate that interactions between nutrients are not universal, but rather they are conserved at the family-level.

## Discussion

Our findings pose intriguing questions about the mechanisms behind the nutrient interaction patterns we have observed. For instance, is it reasonable to expect that the additive null model should have worked as well as it did, and better at the family than at the species level? Why are pairs of more similar nutrients better explained by the null model than pairs of more dissimilar nutrients? What may explain why nutrients dominate over others at the family-level? And why do sugars generally dominate organic acids for most families?

We have previously shown that many of the properties of our experimental enrichment communities reflect the generic emergent behavior of consumer-resource models [14,17], and subsequent work extended this finding to complex natural communities [18]. We thus sought to ask whether our observations regarding the assembly of communities in pairs of resources are similarly reflecting a generic emergent property of consumer-resource models. To address this question, we followed the same procedure as we and others have done in previous work [14,18–20], and simulated the top-down assembly of microbial communities in pairs *vs* single nutrients using a recently developed Microbial Consumer Resource Model (MiCRM) [14,17,18] (see Methods). The MiCRM differs from the classical MacArthur-Levins model [21] in that it includes metabolic cross-feeding in a manner that preserves thermodynamic balance. The model and the details of the simulations are described in the Methods section. In brief, 200 species are seeded into each habitat at the start of a simulation. Each of these is represented by a different vector of resource uptake rates. These vectors are randomly sampled in a manner that captures the existence of two functional guilds, each of which specializes in a different group of resources (e.g. sugar vs organic acid specialists) (**Fig. 4A**). We note that this specialization is quantitative rather than discrete, as all species are assumed to be able to consume all of the resources (a point that is in general consistent with our experimental findings (**Fig. S9**)). Communities are allowed to find a dynamical equilibrium, at which point we stop the simulation. In total, and in order to get to generic behavior, we generated 100 simulations each with a different random set of species (Methods).

**Fig. 4.**
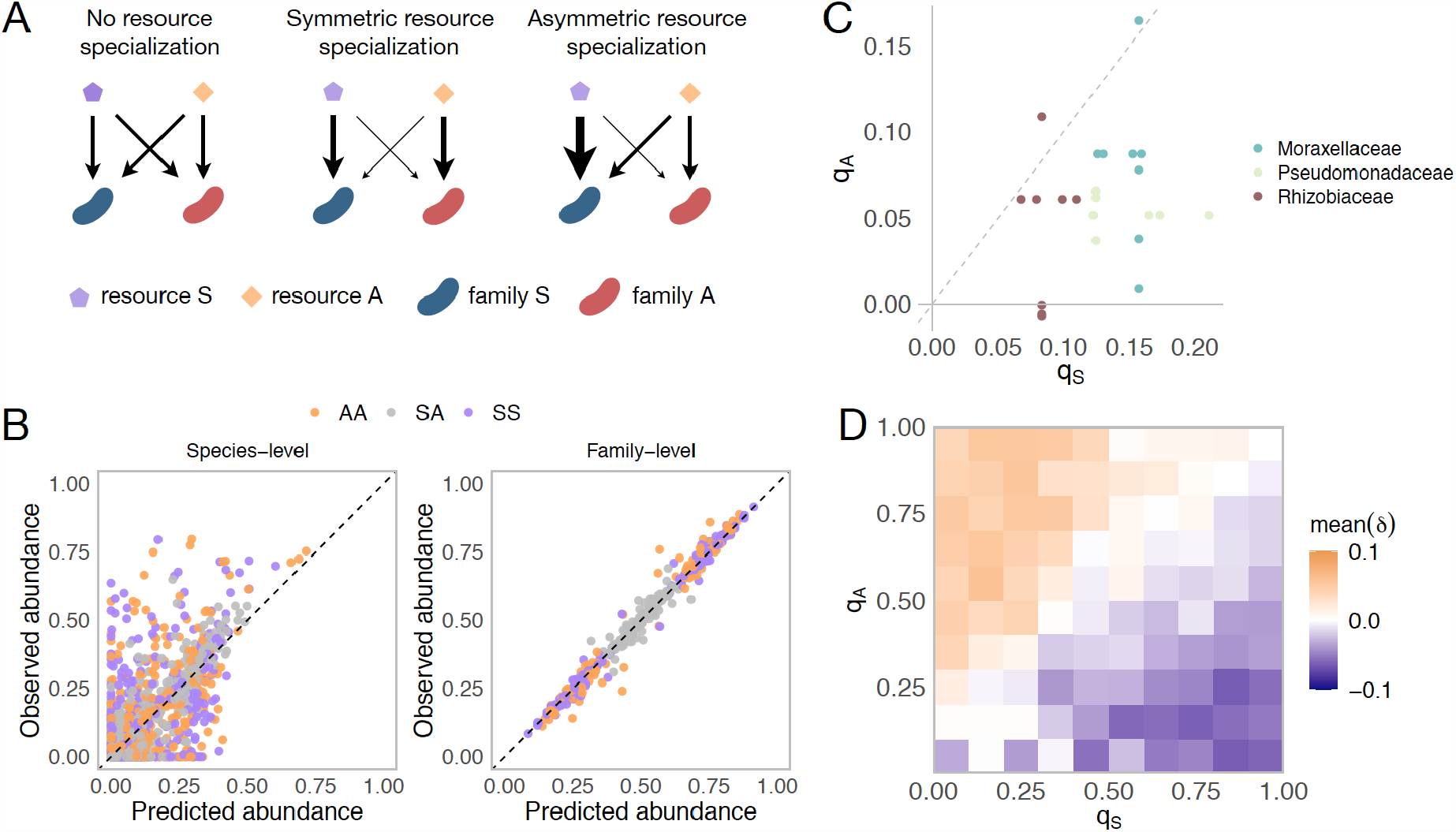
Family-level asymmetry in nutrient benefits can lead to dominance. **(A)** Schematic illustrating different scenarios of nutrient preference. There are two families (*F*_*S*_ and *F*_*A*_) and two resource classes (*R*_*S*_ and *R*_*A*_). Without resource specialization, *F*_*S*_ and *F*_*A*_ have equal access to *R*_*S*_ and *R*_*A*_. With symmetric specialization, each family prefers its own resource class with the same strength. With asymmetric specialization, one family (*F*_*S*_) has better access to its own resource class (*R*_*S*_) relative to that of the other family (*F*_*A*_) on its own resource class (*R*_*A*_). (**B)** A mechanistically-explicit consumer-resource model that incorporates resource competition, resource specialization and nonspecific cross-feeding (Methods) recovers the predicted additivity pattern at both the species (left) and family (right) level of taxonomic organization. The observed relative abundance of each species or family in 300 communities grown on a different pair of nutrients (100 AA, 100 SS, and 100 SA) is plotted against the abundance predicted from the same communities grown on each of the relevant single nutrients (S, A). Each family specializes equally on its preferred nutrient (*q*_*S*_ = *q*_*A*_ = 0.9) as in previous work [18]. In **Fig. S9**, we illustrate representative consumption matrices for different choices of *q*_*A*_ and *q*_*S*_. **(C)** 22 strains were isolated from the assembled communities and their growth rates on minimal M9 media supplemented with one the 10 carbon sources were measured. *q*_*S*_ represents the growth rate advantage of Enterobacteriaceae on sugars relative to the other dominant family (coloured) while *q*_*A*_ represents the growth rate advantage of the other family on the acids relative to Enterobacteriaceae (Methods). When *q*_*S*_ is positive (negative), Enterobacteriaceae grow faster (slower) on the sugar than the other family. When *q*_*A*_ is positive (negative), the other family grows faster on the acid than Enterobacteriaceae. Each dot corresponds to a sugar-acid pair for a Enterobacteriaceae-other family pair (n=24). The growth rate advantage of Enterobacteriaceae on sugars is significantly greater than the growth rate advantage of the other families on acids (i.e. *q*_*S*_ >*q*_*A*_, mean of differences = 0.069, paired t-test, n=24, p-value<0.0001). **(D)** Here we repeat the same simulation as shown in **B**, this time using different combinations of *q*_*A*_ and *q*_*S*_ (0.05, 0.15, 0.25, 0.35, 0.45, 0.55, 0.65, 0.75, 0.85, 0.95). Heatmap shows the mean dominance level (*δ*) for different combinations of *q*_*A*_ and *q*_*S*_. When *δ* <0, the sugar dominates (purple); when *δ* >0, the acid dominates (orange).

A generic property of these simulations is that the species-level community composition on mixtures of two limiting nutrients is reasonably well described by the additive null model (Pearson’s R=0.7 and p<0.001; RMSE=0.097; N=2440) (**Fig. 4B**; Methods), which is consistent with previous consumer-resource modelling work [10,18]. In addition, when we group species by the functional groups they belong to (i.e. family), the predictive ability of the additive null model improves (Pearson’s R=0.99 and p<0.001; RMSE=0.03; N=414) (**Fig. 4B**), a point that is consistent with our experimental findings (**Fig. S4**). This family-level additivity holds when communities are randomly colonized by a different set of species (**Fig. S10**), suggesting that family-level additivity is robust to species-level taxonomic variability.

By contrast, the simulated communities do not exhibit any systematic dominance, neither at the species nor at the family level (**Fig. 4B**). What feature of the MiCRM might be causing us to miss this experimental behavior? One assumption of the model, which we had made for the sake of simplicity and for consistency with previous work, is that all nutrients are equally valuable for the microbes that specialize on them. In other words, the benefits of growing in each type of nutrient are symmetric. Yet, this assumption is not really consistent with the empirical reality that glucose specialists, such as Enterobacteriaceae, grow more strongly in sugars than organic acid specialists do on organic acids. This is illustrated in **Fig. 4C**, where we plot the growth advantage for 7 Enterobacteriaceae isolates in sugar media *vs* the growth advantage of Pseudomonadaceae, Rhizobiaceae and Moraxellaceae isolates in organic acids.

We postulated that including this asymmetry may unbalance the competition for resources and give rise to nutrient dominance at the family-level, as the family that lies on the winning side of that asymmetry may leverage its enhanced competitive ability in the most valuable nutrient to displace the losing family from its lower-value nutrient niche. To test this intuition, we relaxed the symmetry in resource value that was imposed by default in the model, and repeated our simulations for different levels of nutrient value asymmetry (the simulations still include facilitation via metabolite secretion, as we had done in all prior simulations) (**Fig. 4A, Fig. S11**). As we show in **Fig. 4D**, and consistent with our intuition, nutrient dominance at the family-level may emerge as a generic property of microbial consumer-resource models when a nutrient is substantially more valuable than the other. Reassuringly, our experiments indicate that dominance is generally favorable to the taxa that benefits from growth asymmetry e.g. to Enterobacteriaceae in sugar-acid mixes, and unfavorable to families in the losing end of growth asymmetry (Pseudomonadaceae, Rhizobiaceae and Moraxellaceae) in those same environments. This observation is consistent with the behavior of the model (**Fig. 4D**).

In sum, our analysis indicates that our empirical observations regarding the assembly of microbial communities in nutrient mixes are consistent with generic behavior of consumer-resource models. Based on this finding, we cautiously suggest that family-level asymmetries in nutrient benefits may be a possible mechanism for the general nutrient dominance patterns we have observed, and that a null additive model is in general a good first approximation for the assembly of microbial communities in simple nutrient mixtures (a pattern that is consistent with previous work [9,10]). It is important to recognize, however, that other explanations and mechanisms may be at play too. Quantitatively elucidating the specific mechanisms that may explain the individual patterns of nutrient interactions (or lack thereof) for each family and in each pair of nutrients would require us to measure the amounts of all nutrients secreted by every species in each environment over time (i.e. in each nutrient and in each pair) and then characterize the growth curves of all species in those nutrients. Although such monumental effort is beyond the scope of this paper, we hope that our findings and methodology will be a stepping stone towards elucidating how microbial communities assemble in complex nutrient mixtures, and that they will stimulate further theoretical and empirical work. We propose that top-down community assembly in combinatorially reconstructed nutrient environments can be a helpful approach not only to understand the origins of microbial biodiversity, but also to learn how to manipulate existing microbiomes by rationally modulating nutrient availability.

## Material and Methods

### Null model for relative abundance

Let’s consider a simple scenario of two co-cultures of species A and B growing together in two separate demes, each containing a single nutrient (labeled 1 and 2). The fractions of A and B in nutrient/deme 1 are *f*_*A,1*_ = *n*_*A,1*_ /(*n*_*A,1*_+*n*_*B,1*_) and *f*_*B,1*_ = *n*_*B,1*_/(*n*_*A,1*_+*n*_*B,1*_) respectively, and similarly, the fractions of A and B in nutrient/deme 2 are *f*_*A,2*_ = *n*_*A,2*_/(*n*_*A,2*_ +*n*_*B,2*_) and *f*_*B,2*_ = *n*_*B,2*_ /(*n*_*A,2*_+*n*_*B,2*_) (where *n* is the total number of cells of species A or B). If we consider the two-deme system as a whole (i.e. if we pool together the amount of species in each nutrient/deme) the fractions of A and B in the mixture are given by:

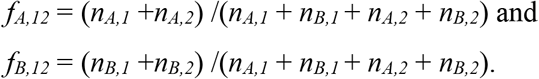

We can define *n*_*t,1*_ = *n*_*A,1*_+*n*_*B,1*_ and *n*_*t,2*_ = *n*_*A,2*_+*n*_*B,2*_ as the total number of cells in the nutrient demes 1 and 2, respectively. We can thus write *f*_*A,12*_ = (*n*_*A,1*_+*n*_*A,2*_) /(*n*_*t,1*_+*n*_*t,2*_). Defining *w*_*1*_ = *n*_*t,1*_/(*n*_*t,1*_+*n*_*t,2*_) and *w*_*2*_ = *n*_*t,2*_/(*n*_*t,1*_+*n*_*t,2*_), it is straightforward to show that: *f*_*A,12*_ = *w*_*1*_ *f*_*A,1*_ + *w*_*2*_ *f*_*A,2*_. By the same reasoning, we find that *f*_*B,12*_ = *w*_*1*_ *f*_*B,1*_ + *w*_*2*_ *f*_*B,2*_.

### Sample collection

Soil samples were collected from two different natural sites in West Haven (CT, USA), with sterilized equipment, and placed into sterile bottles. Once in the lab, five grams of each soil sample were then transferred to 250mL flasks and soaked into 50mL of sterile 1xPBS (phosphate buffer saline) supplemented with 200 μg/mL cycloheximide (Sigma, C7698) to inhibit eukaryotic growth. The soil suspension was well mixed and allowed to sit for 48h at room temperature. After 48h, samples of the supernatant solution containing the ‘source’ soil microbiome were used as inocula for the experiment or stored at −80°C with 40% glycerol.

### Preparation of media plates

Carbon source stock solutions (CS, **Table S1**) were prepared at 0.7 C-mol/L (10x) and sterilized through 0.22µm filters (Millipore). CS were aliquoted into 96 deep-well plates (VWR) as single CS or mixed in pairs at 1:1 (vol:vol) and stored at -20°C. To keep the total amount of carbon constant across all treatments, pairs contained half the amount of each carbon source compared to their respective single CS. Synthetic minimal growth media was prepared from concentrated stocks of M9 salts, MgSO_4_, CaCl_2_, and 0.07C-mol/L (final concentration) of single or pairs of CS. The final pH of all growth media is shown in **Table S1**.

### Community assembly experiment

Starting inocula were obtained directly from the ‘source’ soil microbiome solution by inoculating 40μL into 500μL culture media prepared as indicated above. For each sample and CS, 4 μL of the culture medium was dispensed into fresh media plates containing the different single or pairs of CS in quadruplicate. Bacterial cultures were allowed to grow for 48h at 30°C in static broth in 96 deep-well plates (VWR). After 48h each culture was homogenized by pipetting up and down 10 times before transferring 4 μL to 500μL of fresh media, and cells were allowed to grow again. Cultures were passaged 10 times (∼70 generations). OD620 was measured after 48h growth. Samples were frozen at -80°C with 40% glycerol.

### DNA extraction, library preparation, and sequencing

Samples were centrifuged for 40mins at 3500rpm, and the pellet was stored at -80°C until DNA extraction. DNA extraction was performed with the DNeasy 96 Blood & Tissue kit for animal tissues (QIAGEN), as described in the kit protocol, including the pre-treatment step for Gram-positive bacteria. DNA concentration was quantified using the Quan-iTPicoGreen dsDNA Assay kit (Molecular Probes, Inc) and the samples were normalized to 5ng/uL before sequencing. The 16S rRNA gene amplicon library preparation and sequencing were performed by Microbiome Insights, Vancouver, Canada (www.microbiomeinsights.com). For the library preparation, PCR was done with dual-barcoded primers [22] targeting the 16S V4 region and the PCR reactions were cleaned up and normalized using the high-throughput SequalPrep 96-well Plate Kit. Samples were sequenced on the Illumina MiSeq using the 300-bp paired end kit v3.chemistry.

### Taxonomy assignment

The taxonomy assignment was performed as described in previous work [15]. Following sequencing, the raw sequencing reads were processed, including demultiplexing and removing the barcodes, indexes and primers, using QIIME (version1.9, [23]), generating fastq files with the forward and reverse reads. DADA2 (version 1.6.0) was then used to infer exact sequence variants (ESVs) [24]. Briefly, the forward and reverse reads were trimmed at position 240 and 160, respectively, and then merged with a minimum overlap of 100bp. All other parameters were set to the DADA2 default values. Chimeras were removed using the “consensus” method in DADA2. The taxonomy of each 16S exact sequence variant (ESV) was then assigned using the naïve Bayesian classifier method [25] and the Silva reference database version [26] as described in DADA2. The analysis was performed on samples rarefied to 10779 reads.

### Quantification of total abundances, interactions, and dominance

We used OD620 after the 48h growth cycle as a proxy for total population size (community biomass). The predicted relative abundance of species *i* in a mix of nutrients 1 and 2 was then calculated as *f*_*i,12*_(null) = *w*_*1*_ *f*_*i,1*_ + *w*_*2*_ *f*_*i,2*_ where *f*_*i,1*_ and *f*_*i,2*_ are the relative abundances of *i* in nutrients 1 and 2, respectively, and *w*_*1*_ = (OD620_1_/(OD620_1_+OD620_2_) and *w*_*2*_ = (OD620_2_/(OD620_1_+OD620_2_). For each carbon source pair and inoculum, *f*_*i,12*_(null) is calculated as the mean of the two single carbon source-replicate pairwise combinations (N=16). In order to quantify interactions, we first determine whether an interaction between nutrients exists for each nutrient pair (nutrient 1 and nutrient 2) and family *i*. An interaction exists when *ε* = *f*_*i,12*_ *-f*_*i,12*_(null) is significantly different from 0 (one-sample Student’s t-test, p<0.05), that is when there is a deviation from the null prediction. Under such condition (i.e. |*ε*|>0), synergy and antagonism (which are forms of super-dominance) occur when *f*_*i,12*_ > *max*(*f*_*i,1*_, *f*_*i,2*_) and *f*_*i,12*_ < *min*(*f*_*i,1*_, *f*_*i,2*_) respectively, while dominance occurs when *min*(*f*_*i,1*_, *f*_*i,2*_) <= *f*_*i,12*_<*=max*(*f*_*i,1*_, *f*_*i,2*_) (Welch two sample t-test, p<0.05). When *ε*>0, the nutrient with greater abundance dominates; when *ε*<0, the nutrient with lower abundance dominates. For visualization purposes, we developed a dominance index (*δ*). The dominance index for the sugar-acid pairs is written as *δ*_*i*_ = -|*ε*_*12*_| when the sugar dominates and as *δ*_*i*_ = |*ε*_*12*_| when the acid dominates. The dominance index for the sugar-sugar and acid-acid pairs is written as *δ*_*i*_ = -|*ε*_*12*_| when the focal carbon source (glucose or succinate) dominates and as *δ*_*i*_ = |*ε*_*12*_| when the additional carbon source dominates. Pearson’s R was calculated using the R function ‘cor.test’ from the ‘stats’ package and the RMSE was calculated using the ‘rmse’ function from the ‘Metrics’ package.

### Isolation of strains

Several communities (transfer 10) from different inocula and carbon sources were plated on chromogenic agar (HiCrome Universal differential Medium, Sigma) and grown for 48h at 30°C. Single colonies exhibiting distinct morphologies and/or colours were picked, streaked a second time on fresh chromogenic agar plates for purity, and grown for 48h at 30°C. A single colony was then picked from each plate and grown into Tryptic Soy Broth (TSB) for 48h at 30°C. The single-strain cultures were stored with 40% glycerol at -80°C. The isolated strains were sent for full-length 16S rRNA Sanger sequencing (Genewiz) and their taxonomy was assigned using the online RDP naïve Bayesian rRNA classifier version 2.11.

### Growth rate estimation

22 isolated strains belonging to the four dominant families, namely Enterobacteriaceae (7), Pseudomonadaceae (5), Moraxellaceae (6) and Rhizobiaceae (4) (**Table S2**) were streaked from frozen stock on chromogenic agar plates and grown for 48h at 30°C. For each strain, a single colony was pre-cultured in 500ul TSB in a deepwell plate for 24h at 30°C. Each strain was then acclimated into the 10 single carbon sources (glucose, fructose, cellobiose, ribose, glycerol, succinate, fumarate, benzoate, glutamine, and glycine). For this, 2ul of the grown pre-culture was inoculated into 500ul of fresh minimal media with each carbon source at a concentration of 0.07 C-mol/liter and grown for 48h at 30°C. The growth curve assay was then performed in a 384 wellplate by inoculating 1ul of the grown isolate culture on 100ul of fresh media of the same carbon source as for the acclimation step (3-4 replicates each). OD620 was read every 10 min for ∼40h at 30°C. The average growth rate of each strain in each carbon source was calculated as *r*_avg_ = log_2_(*N*_*f*_ /*N*_*i*_)/(*t*_*f*_ -*t*_*i*_) where *N*_*f*_ is the OD at 16h (ie. *t*_*f*_) and *N*_*i*_ is the OD at 0.5h (i.e. *t*_*f*_).

### Growth rate asymmetry calculation

The growth rate asymmetry on sugars (*q*_*S*_) is calculated as *q*_*S*_ = *r*_avg_(*E, S*) - *r*_avg_(*O, S*) where *r*_avg_(*E, S*) is the mean average growth rate of Enterobacteriaceae on the sugar *S* and *r*_avg_(*O, S*) is the mean average growth rate of one of the other dominant families (i.e. Pseudomonadaceae, Moraxellaceae, or Rhizobiaceae) on *S*. The growth rate asymmetry on organic acids (*q*_*A*_) is calculated as *q*_*A*_ = *r*_avg_(*O, A*) - *r*_avg_(*E, A*) where *r*_avg_(*O, A*) is the mean average growth rate of one of the other dominant families (i.e. Pseudomonadaceae, Moraxellaceae, or Rhizobiaceae) on the organic acid *A* and *r*_avg_(*E, A*) is the mean average growth rate of Enterobacteriaceae on *A*.

### Microbial Consumer Resource Model

Microbial community assembly is modelled using the Microbial Consumer Resource Model (MiCRM), with simulations implemented using *Community Simulator*, a freely available Python package [19]. This model has been outlined extensively in previous work and has been shown to qualitatively reproduce ecological patterns across both natural [18] and laboratory [14] microbiomes. Here we describe the exact equations simulated and parameters used in this paper. A more general description of this model is given elsewhere [17,19]. Our MiCRM simulations model the abundance *N*_*i*_ of *n* species and the abundance *R*_*α*_ of *M* resources in a well-mixed chemostat-like ecosystem with continuous resource flow. We focus on continuous resource flow for simplicity and because previous work has shown that the major qualitative features of the MiCRM are unaffected by periodic resource supply (as was the case in our experiments) [14]. Species interact by uptake and release of resources into their environment. The dynamics of the system are governed by the following set of ordinary differential equations:

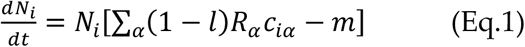

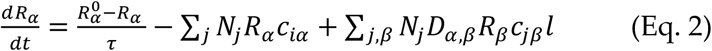

Here *c*_*iα*_ is the uptake rate of resource *α* by species *i, m* is the minimal energy requirement for maintenance of species *i,τ* is the timescale for supply of external resources, 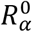 is the abundance of resource *α* supplied (i.e. the abundance in the media), *l* is the fraction of resource secreted as by-product and *D*_*α,β*_ is the fraction of resource *α* secreted as by-product *β*. In line with previous work, the following parameters are kept constant for all simulations *τ* = 1, *m* = 1 and *l* = 0.5 [14,18].

In the MiCRM, by-product production is encoded in the metabolic matrix *D* where each element *D*_*α,β*_ specifies the fraction of resource *α* secreted as by-product *β*. As in previous work, each column *β* in *D*_*α,β*_ is sampled from a Dirichlet distribution with concentration parameter *d*_*α,β*_ = 1/*M*_*s*_ where *s =* 0.3 is a parameter that tunes the sparsity of the underlying metabolic network. The Dirichlet distribution ensures that each column sums to 1 so that the total secretion flux does not exceed the input flux. For simplicity we used a fixed concentration parameter and so are not assuming any underlying metabolic structure. The MiCRM also assumes that all species have the same *D* matrix, i.e when growing on the same resource each species releases the same metabolic by-products.

In our simulations, species differ solely in the uptake rate for different resources *c*_*iα*_ where *i* is the species and *α* is the resource. Taxonomic specialization is introduced in the form of two families *F*_*A*_ and *F*_*S*_ that each have a preference for one of two resource classes *A* and *S*, respectively. Each *c*_*iα*_ is sampled from a gamma distribution (to ensure positivity) whose mean <*c*_*iα*_> and variance var(*c*_*iα*_) depends on the family of *i* and the resource class of *α*. This means that all species are capable of metabolizing all resources. Specifically

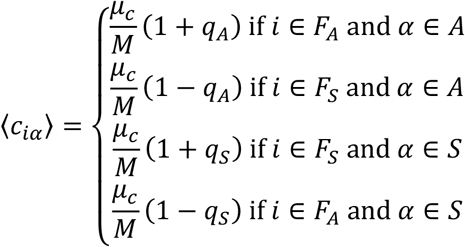

and

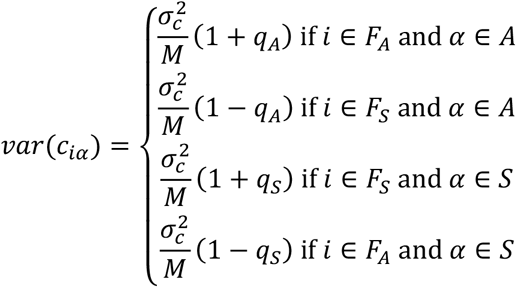

*μ*_*c*_ = 10 determines the overall mean uptake rate and 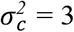 determines overall variance in uptake rate (these parameters are the default value in the *Community Simulator* package). Parameters *q*_*A*_ and *q*_*A*_ tune the relative advantage each specialist family has on its preferred resource. When *q*_*A*_ = 1, only *F*_*A*_ consumes resources in *A* whereas when *q*_*A*_ = 0 both families have equal access to resources in *A*. Conversely when *q*_*S*_ = 1, only *F*_*S*_ consumes resources in *S* whereas when *q*_*S*_ = 0 both families have equal access to resources in *S*.

For each simulation we consider 200 species (100 per family). Each community in one simulation is seeded with all 200 species. This means that there is no stochasticity in colonisation (though see **Fig. S10** where this assumption is relaxed). We choose 200 species as this is within the range of the number of ESVs in a typical inoculum for our experiments (110-1290 ESVs, reported in [14]). The initial abundances are all set to 1 for simplicity. In line with our experiments, either one or two resources are supplied in the media and the rest are generated as metabolic by-products. For simulations with a single supplied resource, 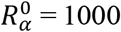 if *α* is the supplied resource and 0 otherwise. For simulations with two supplied resources, 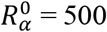 for each supplied resource and 0 otherwise. This ensures that the total amount of resources is kept constant as in our experiments. In total we consider 20 resources in each simulation (with 10 resources in each resource class (A or S)) as this gives us communities with 7±2 species (Mean±SD) at equilibrium, which is comparable to the diversity of our experimental communities.

In line with our experiments, each simulation consisted of three types of mixed resource environments (one with two supplied resources in class *R*_*A*_, one with two supplied resources in class *R*_*S*_ and one with one resource in class *R*_*A*_ and one resource in class *R*_*S*_). We also included all 4 single resource environments needed to predict the mixes (i.e. two with the resources in class *R*_*A*_ and two with the resources in class *R*_*s*_). Therefore, each simulation consisted of 7 communities each in a different environment and all seeded with the same initial set of 200 species. The equilibrium for all 7 communities was found using the `SteadyState` function in *Community Simulator* [19]. Failed runs where the `SteadyState` function returned an error were removed from our analysis. In addition, for each simulation we tested that the `SteadyState` algorithm had truly converged to an equilibrium using the same approach as in [18] and removed all non-convergent runs (defined as a run for which | *d ln*(*N*_*i*_) / *dt* | > *10*^*−5*^). Including these runs would not have qualitatively changed our results.

In the raw numerical output of the run, all species have non-zero abundances due to limits in numerical precision. A species was considered extinct if its abundance was less than 10^−6^ which was set by looking at a histogram of the raw output of our simulations. Once the extinct species were removed, we predicted the relative abundance of each species *i* in the mixture of nutrients using the same approach that had been used for the experimental data. To obtain a statistically robust sample size, we repeated this procedure for 100 replicate simulations, resampling all randomly generated parameters in each simulation (i.e. resampling all *c*_*iα*_ and *D*_*α,β*_ as described above).

## Acknowledgments

We want to thank members of the Sanchez lab for helpful discussions. This work was supported by the National Institutes of Health through grant 1R35 GM133467-01, and by a Packard Foundation Fellowship to AS.

## Competing interests

The authors declare no competing interests.

## Notes

### Competing Interest Statement

The authors have declared no competing interest.

