## Supplementary Material for "Nutrient dominance governs the assembly of microbial communities in mixed nutrient environments"

This pdf includes:
Figures S1 to S11
Tables S1 and S2

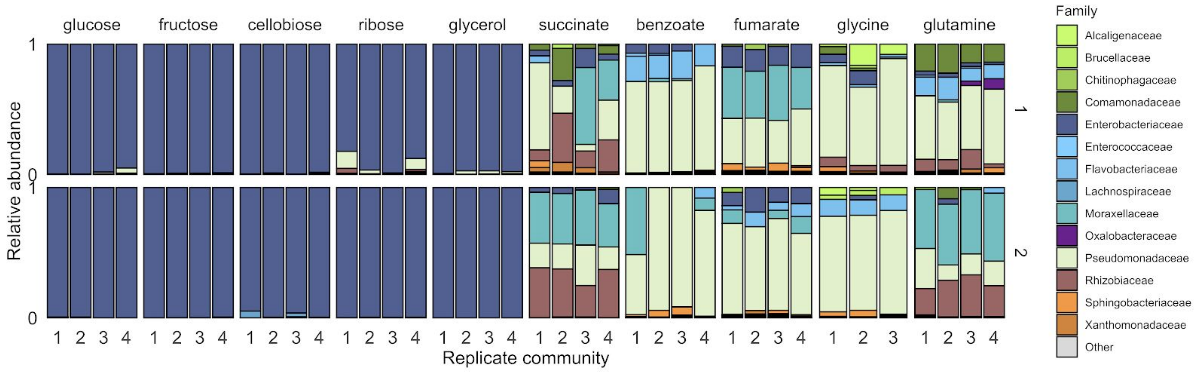


**Fig. S1. Community assembly in a single carbon source.** Two soil samples were inoculated in minimal M9 medium supplemented with a single carbon source (three or four replicates each), and propagated into fresh media every 48h for 10 transfers (Methods). Shown is the family-level taxonomic composition at Transfer 10 for inoculum 1 (top) and inoculum 2 (bottom). Families with a relative abundance lower than 0.01 are shown as ‘Other’.

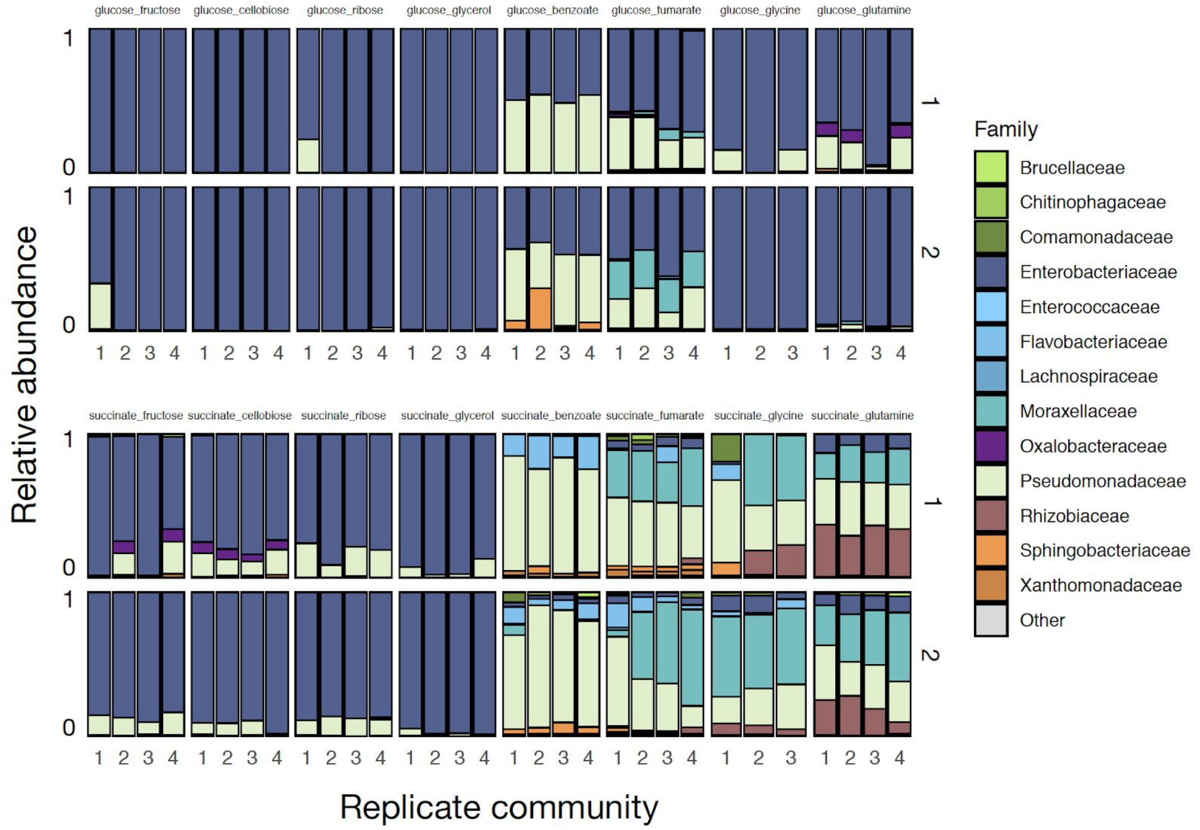


**Fig. S2. Community assembly in a mixture of two carbon sources.** Two soil samples were inoculated in minimal M9 medium supplemented with two carbon sources (glucose or succinate + another carbon source), and propagated into fresh media every 48h for 10 transfers (Methods). There are three/four replicates per carbon source pair. Shown is the family-level taxonomic composition at Transfer 10 for the two inocula. Families with a relative abundance lower than 0.01 are shown as ‘Other’.

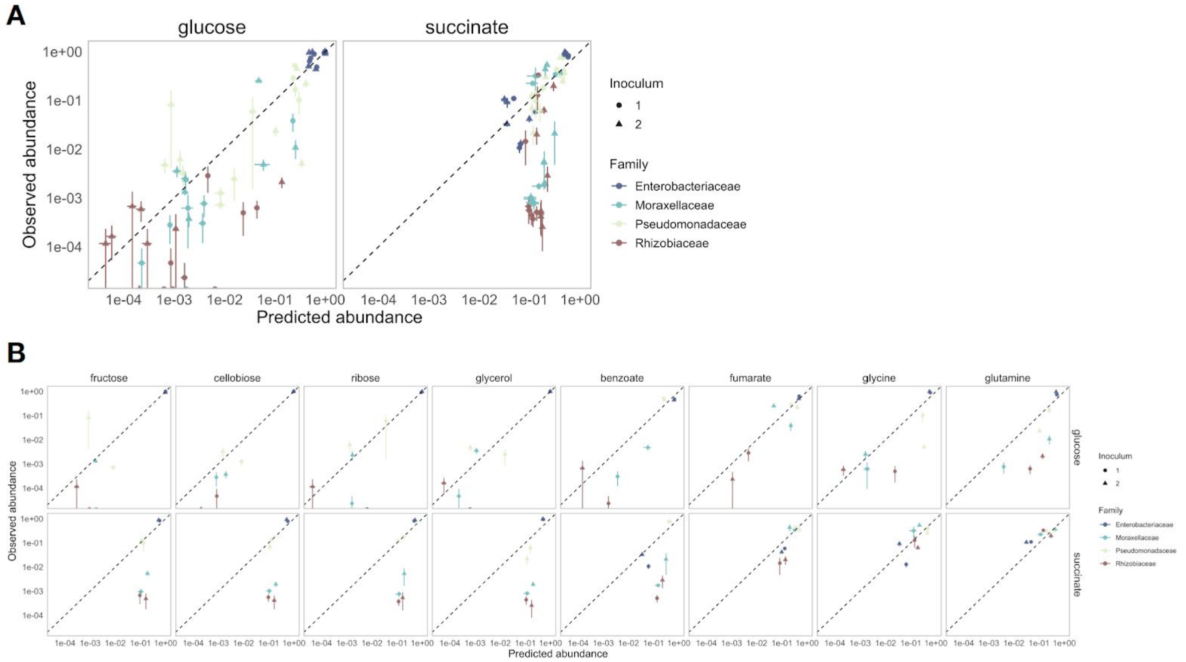


**Fig. S3. Systematic deviations from the null (additive) prediction reveal interactions between nutrients.** Shown is the same data as in **Fig. 2B** (**A**) and **Fig. 2C** (**B**) but displayed on a log-log scale so the datapoints at lower relative abundance are easier to visualise.

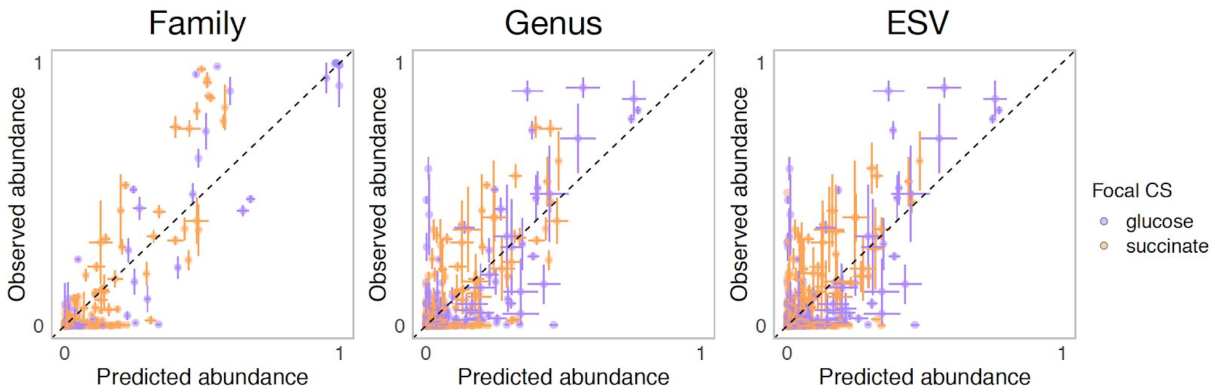


| Level | Focal CS | Pearson's R | RMSE |
|---|---|---|---|
| Family | glucose | 0.948 | 0.073 |
| Family | succinate | 0.871 | 0.094 |
| Genus | glucose | 0.782 | 0.082 |
| Genus | succinate | 0.765 | 0.066 |
| ESV | glucose | 0.773 | 0.069 |
| ESV | succinate | 0.674 | 0.056 |

**Fig. S4. Comparison of the observed relative abundance and abundance predicted by the null model.** Shown is the observed vs predicted abundance for different taxonomic levels and focal carbon source (CS) (mean ± SE). Table shows the Pearson's R and RMSE for each family-focal carbon source combination.

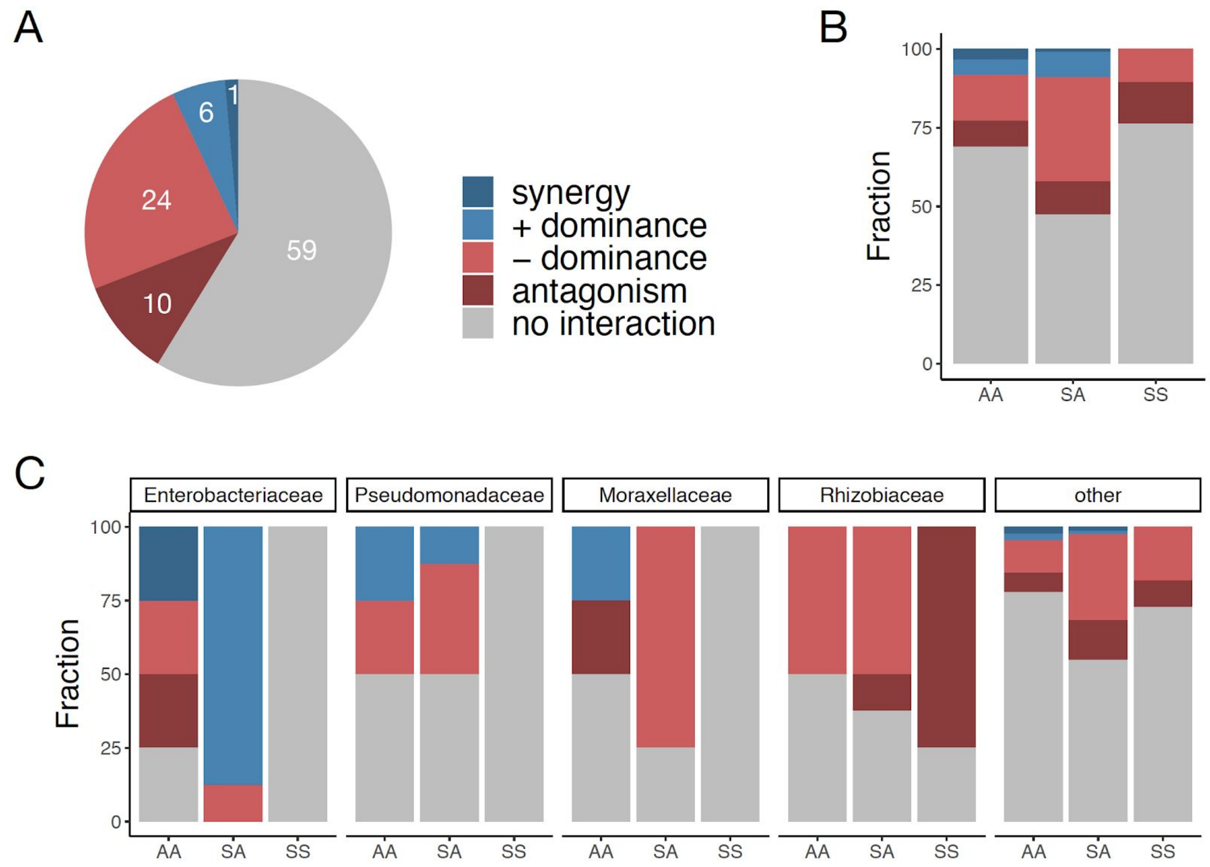


**Fig. S5. Dominance is the most common type of nutrient interaction, especially in the sugar-acid mixtures. (A)** Interaction type for each pair of carbon source and family. An interaction between nutrients occurs when $\varepsilon$ is significantly different from 0 for the two inocula x four replicates (one-sample Student's t-test, $p < 0.05$) (Methods). Multiple types of interaction are possible: dominance, synergy, and antagonism. Synergy (antagonism) occurs when the abundance in the mixture is greater (lower) than the abundances in any of the single nutrients independently (Welch two sample t-test, $p<0.05$) (Methods). Dominance occurs when the abundance in the mixture is closer or similar to the abundance in one of the singles. **(B)** Interaction type by carbon source pair type. AA: mixture of two acids; SS: mixture of two sugars; and SA: mixture of a sugar and an acid. **(C)** Interaction type shown for the four most abundant families and 'other' families grouped together.

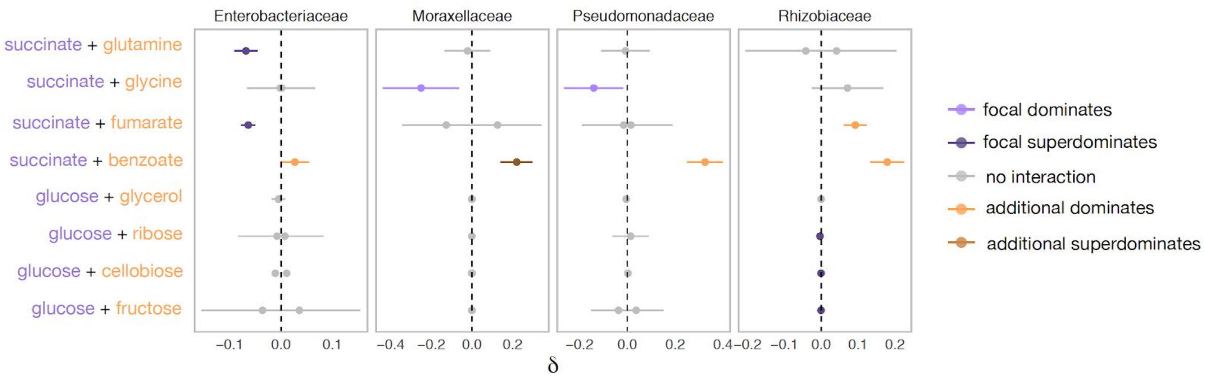


**Fig S6. Family-level dominance for mixtures of acid-acid and sugar-sugar.** When datapoints lie on the left of the 0 line, the focal carbon source (succinate or glucose) dominates. When datapoints lie on the right of the 0 line, the additional carbon source dominates (Methods). Gray means that there is no interaction, that is, $\varepsilon$ is significantly not different from 0 (one-sample Student's t-test, p-value<0.05, N=8), and thus $\delta$~0, or that dominance is undefined because one carbon source dominates in one of the inocula and the paired carbon source dominates in the other inocula (in which case $\delta$ is shown as both -$\delta$ *and* +$\delta$). Lighter orange or purple indicates dominance while darker orange or purple indicates super-dominance (synergy or antagonism) (Methods). Only the four most dominant families are shown. Mean ± SD.

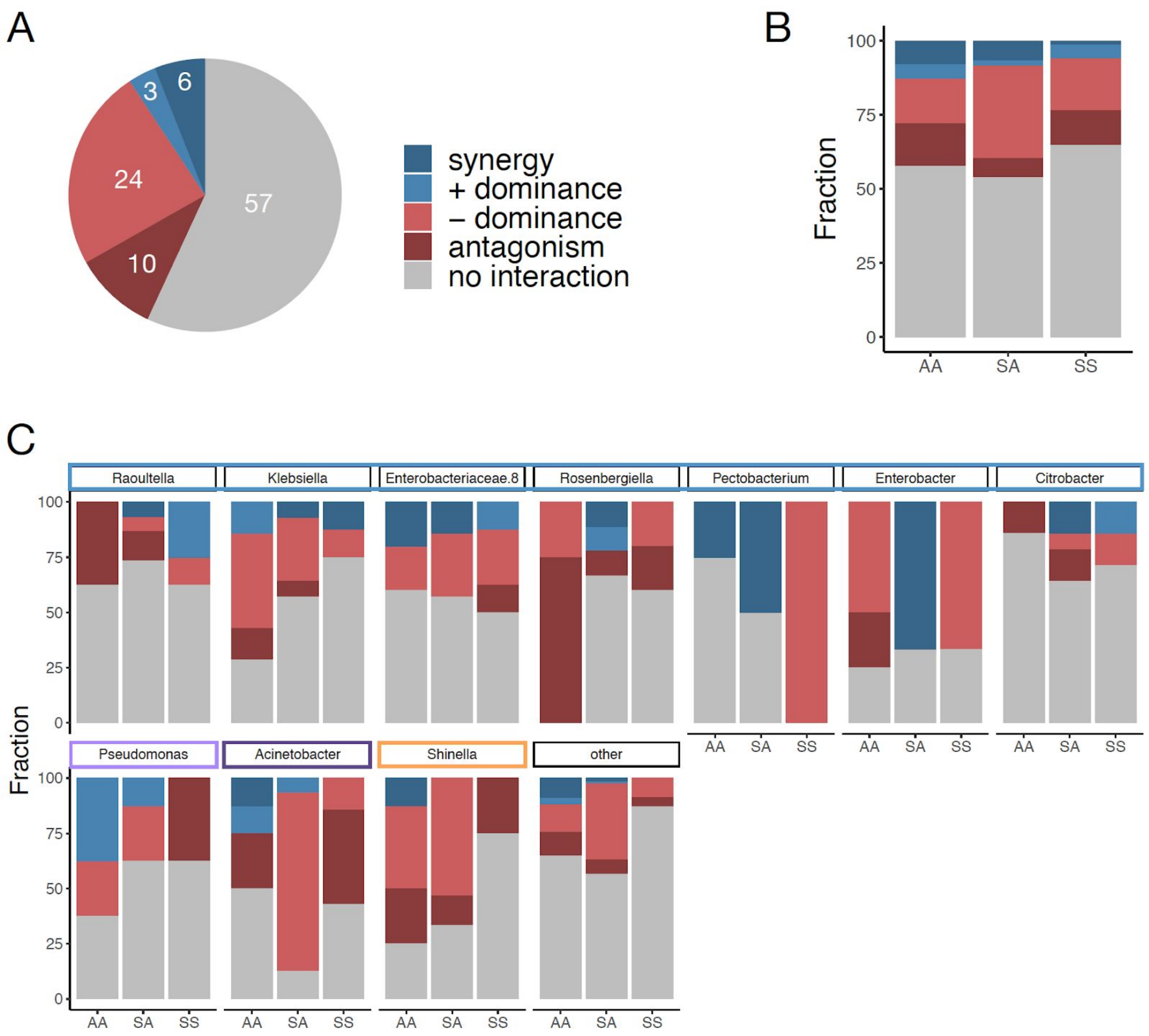


**Fig. S7. Patterns of nutrient interaction at the Genus-level. (A)** Multiple types of nutrient interactions are possible, including dominance, synergy and antagonism (**Fig. 3A**). Interactions occur when $\varepsilon$ is significantly different from 0. Synergy (antagonism) occurs when the abundance in the mixture is greater (lower) than the abundances in any of the singles separately. Dominance occurs when the abundance in the mixture is closer or similar to one of the single abundances but not above or below any of the single abundances independently (Methods). Statistical significance was determined using a t-test ($p < 0.05$) for each pair of carbon source, inocula, and genus. (**B)** Interaction type by carbon source pair type. AA: mixture of two acids; SS: mixture of two sugars; and SA: mixture of a sugar and an acid. (**C)** Interaction type is broken down by the ten most abundant genera spanning the Enterobacteriaceae family (blue), Pseudomonadaceae (light purple), Moraxellaceae (dark purple), and Rhizobiaceae (orange) families. Note that Enterobacteriaceae.8 is a non-identified genus belonging to the Enterobacteriaceae family. The other genera are grouped together and shown as 'other'.

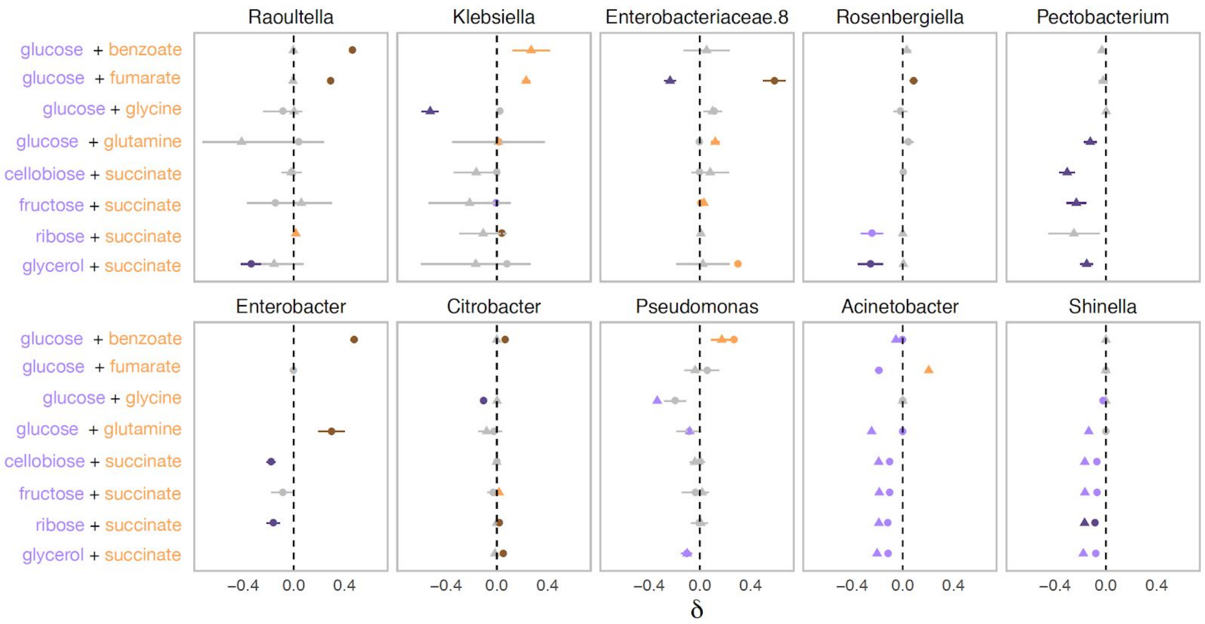


**Fig. S8. The systematic dominance of sugars observed at the family-level does not apply to the genus-level.** To determine the genus-level dominance, the two inocula are considered separately (different shapes) as the genera that are sampled in one inocula may not be sampled in the other inocula. Purple indicates that the sugar dominates while orange indicates that the acid dominates. Lighter purple and orange indicate dominance while darker purple and orange indicate super-dominance (synergy or antagonism) (Methods). No interaction is shown in gray, which occurs when $\varepsilon$ is significantly not different from 0 (one-sample Student's t-test, $p < 0.05$). Shown are the ten most abundant genera (mean ± SD). Note that Enterobacteriaceae.8 is a non-identified genus of the Enterobacteriaceae family.

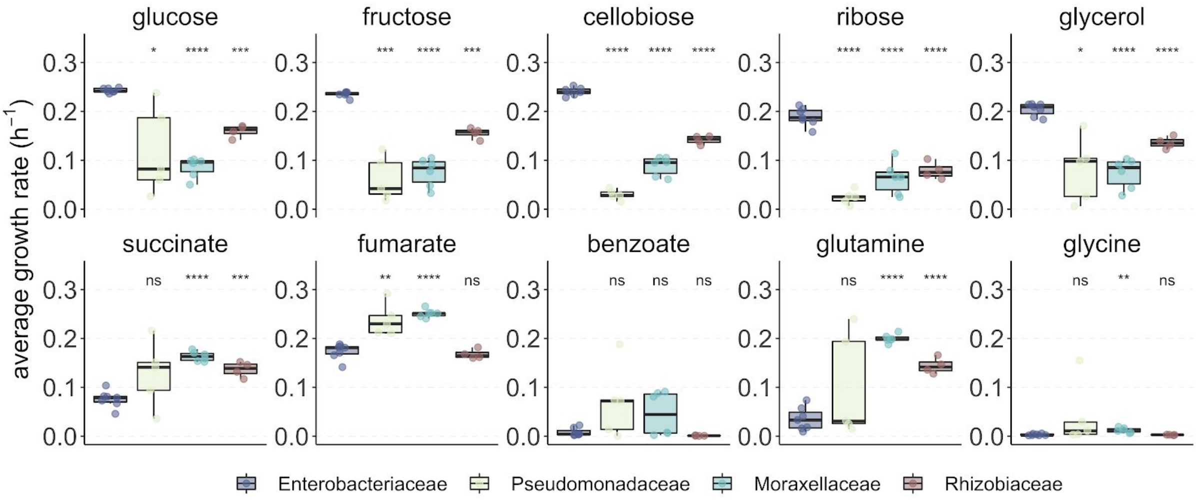


**Fig. S9. Enterobacteriaceae generally have a strong growth advantage in sugars.** 22 strains belonging to the four dominant families, namely Enterobacteriaceae (7), Pseudomonadaceae (5), Moraxellaceae (6) and Rhizobiaceae (4) were isolated from the self-assembled communities and their growth rate on the 10 carbon sources was measured (Methods, **Table S2**). The average growth rate is measured as the mean cell divisions from 0.5h to 16h of growth (3-4 replicates each) (Methods). Significance level (p-value) is measured by comparing the average growth rate between Enterobacteriaceae (reference) and each other family (paired t-test, ****:p<0.0001; ***:p<0.001; **:p<0.01; *:p<0.1).

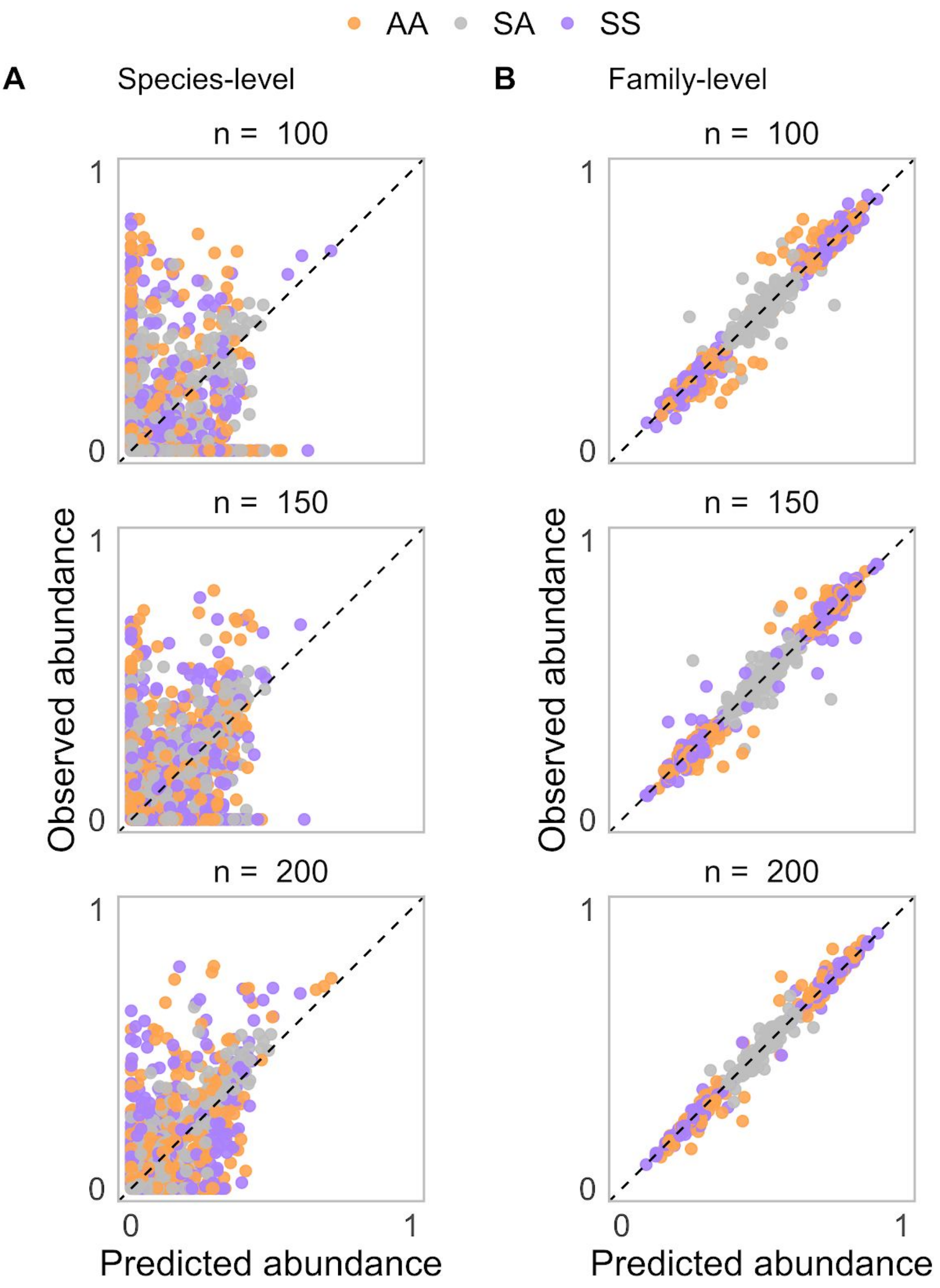


**Fig. S10. Stochastic Colonization has no qualitative effect on the pattern of additivity found using a Microbial Consumer Resource Model.** Relative abundance of each species (**A**) or species grouped by family (**B**) in simulated communities grown in a mixture of nutrients plotted against the predicted relative abundance from simulated communities grown in single nutrients assuming that nutrients act independently (Methods). Communities are colonized with *n* species, randomly sampled from a regional pool of 200 species, while keeping the number of families constant. When *n*=200, all species are sampled. Decreasing *n* reduces the initial species variability of the community and also introduces stochastic colonization through the random sampling of the regional species pool. For each *n*, the result of 100 simulations for communities grown in 3 carbon source pairs is shown (1 SS pair, 1 AA pair and 1 SA pair).

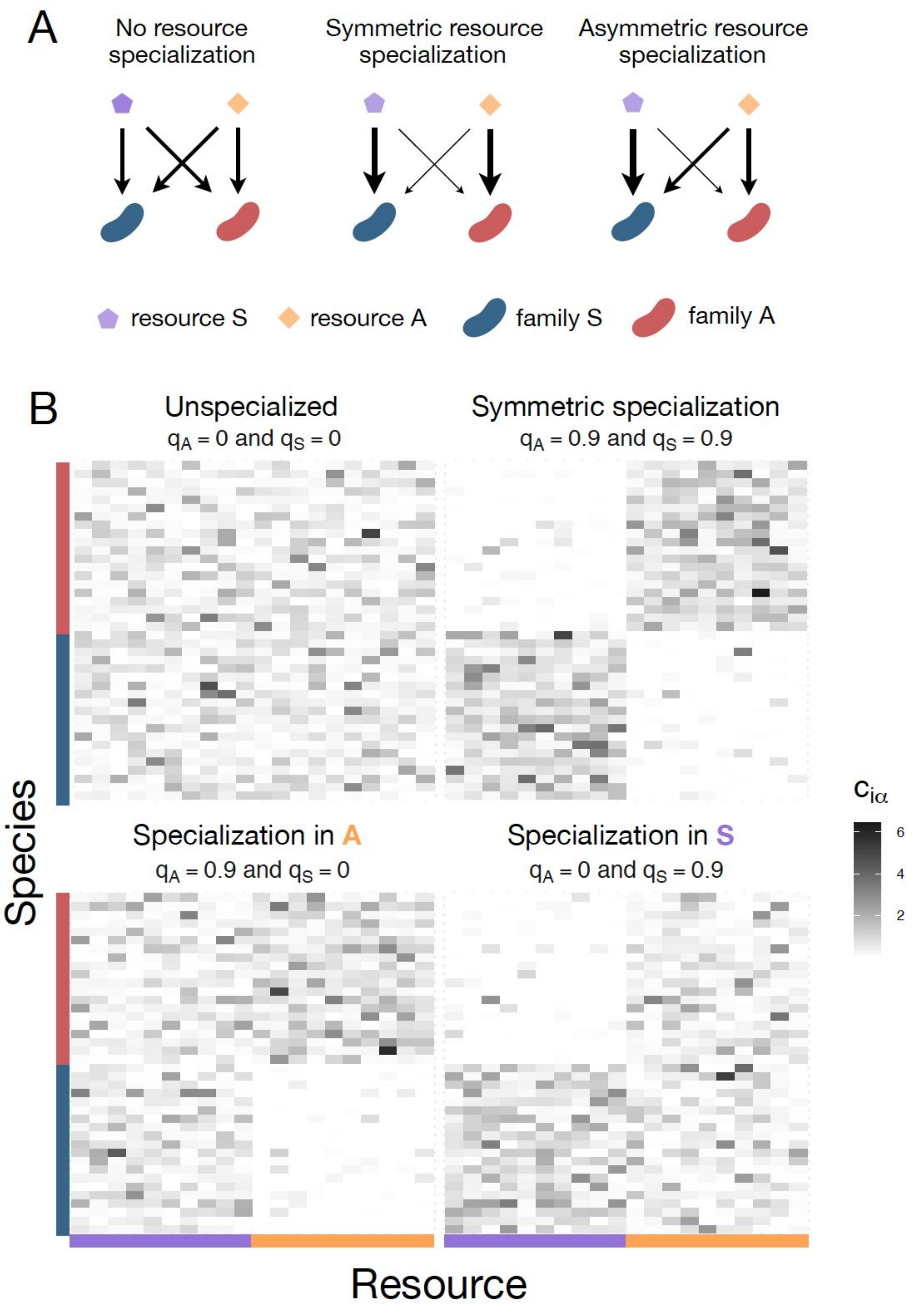


**Fig. S11. Consumption matrices for different patterns of nutrient preference between families used in the consumer-resource model simulations. (A)** The schematics illustrate different scenarios of nutrient preference for two families ($F_S$ and $F_A$) and two resource classes ($R_S$ and $R_A$). Without resource specialization, $F_S$ and $F_A$ have equal access to $R_S$ and $R_A$. With symmetric specialization, each family prefers its own resource class with the same strength. With asymmetric specialization, one family ($F_S$ ) has better access to its own resource class ($R_S$) relative to that of the other family ($F_A$ ) on its own resource class ($R_A$). **(B)** Consumption matrices for two families ($F_A$ and $F_S$ coloured in orange and purple respectively) in two resource classes ($R_S$ and $R_A$). Each row corresponds to a different species (for visualization purposes we show 30 species per family) and each column corresponds to a different nutrient within a resource class (10 nutrients per resource class). The value $c_{i\alpha}$ corresponds to the uptake rate of species $i$ in nutrient $\alpha$. Four nutrient preference patterns are illustrated. Without family-level nutrient preference (specialization), species from the two families have equal access to resources in A ($q_A$=0) and resources in S ( $q_S$ =0). When each family has a strong and quantitatively similar preference for its own resource class, there is symmetric specialization ($q_A = q_S$ >0). When family $F_A$ has a strong preference for its own resource class A but both families have equal access to resources in S, then $q_A$>0 and $q_S$ =0. When family $F_S$ has a strong preference for its own resource class S but both families have equal access to resources in A, then $q_S$>0 and $q_A$=0.

**Table S1. Carbon sources used in this study**

| Carbon source | Supplier | Reference | pH (in M9) |
|---|---|---|---|
| D-Glucose | VWR | 0188-500 | 6.83 |
| D-Cellobiose | Sigma | 22150-10G | 6.84 |
| D-Fructose | Acros Organics | 161355000 | 6.79 |
| D-Ribose | Acros Organics | AC132361000 | 6.81 |
| Glycerol (80%, w/v) | Teknova | G8797 | 6.81 |
| Sodium Succinate hexahydrate | Alfa Aesar | 419A3 | 6.84 |
| Sodium hydrogen fumarate | Alfa Aesar | B24683 | 6.11 |
| Sodium benzoate | Alfa Aesar | A15946 | 6.80 |
| L-Glutamine 200mM (29.23 mg/mL) | Sigma | G7513-100ML | 6.80 |
| Glycine | Sigma | G7126-100G | 6.82 |

**Table S2. Taxonomy of strains used in the growth rate assay and community they were isolated from.**

| Family | Genus | Transfer_CarbonSource_Inoculum_Replicate |
|---|---|---|
| Enterobacteriaceae | Raoultella | T10_glucose_I1_R2 |
| Enterobacteriaceae | Citrobacter | T10_glucose-cellobiose_I1_R1 |
| Enterobacteriaceae | Klebsiella | T10_glucose-cellobiose_I1_R1 |
| Enterobacteriaceae | Citrobacter | T10_succinate_I2_R1 |
| Enterobacteriaceae | Enterobacter | T10_succinate_I2_R1 |
| Enterobacteriaceae | Klebsiella | T10_succinate_I2_R4 |
| Enterobacteriaceae | Raoultella | T10_glutamine_I2_R2 |
| Moraxellaceae | Acinetobacter | T10_succinate_I2_R1 |
| Moraxellaceae | Acinetobacter | T10_succinate_I2_R1 |
| Moraxellaceae | Acinetobacter | T10_succinate_I2_R4 |
| Moraxellaceae | Acinetobacter | T10_succinate_I2_R4 |
| Moraxellaceae | Acinetobacter | T10_glutamine_I2_R2 |
| Moraxellaceae | Acinetobacter | T10_glutamine_I2_R2 |
| Pseudomonadaceae | Pseudomonas | T10_glutamine_I2_R3 |
| Pseudomonadaceae | Pseudomonas | T10_ribose_I1_R1 |
| Pseudomonadaceae | Pseudomonas | T10_benzoate_I1_R3 |
| Pseudomonadaceae | Pseudomonas | T10_fumarate_I2_R2 |
| Pseudomonadaceae | Pseudomonas | T10_benzoate_I2_R3 |
| Rhizobiaceae | Rhizobium | T10_succinate_I2_R1 |
| Rhizobiaceae | Rhizobium | T10_succinate_I2_R1 |
| Rhizobiaceae | Rhizobium | T10_succinate_I2_R4 |
| Rhizobiaceae | Rhizobium | T10_glutamine_I2_R2 |
